# It is the Frequency that Matters --- Effects of Electromagnetic Fields on the Release and Content of Extracellular Vesicles

**DOI:** 10.1101/2023.08.08.552505

**Authors:** Yihua Wang, Gregory A. Worrell, Hai-Long Wang

**Affiliations:** Neurology Department, Mayo Clinic, Rochester, Minnesota; Department of Physiology and Biomedical Engineering, Mayo Clinic, Rochester, Minnesota

## Abstract

Extracellular vesicles (EVs) are small membrane-bound structures that originate from various cell types and carry molecular cargo to influence the behavior of recipient cells. The use of EVs as biomarkers and delivery vehicles for diagnosis and treatment in a wide range of human disease is a rapidly growing field of research and clinical practice. Four years ago, we postulated the hypothesis that electromagnetic fields (EMF) will influence the release and content of EVs (*1*). Since then, we have optimized several technical aspects of our experimental setup. We used a bioreactor system that allows cells to grow in a three-dimensional environment mimicking *in-vivo* conditions. We designed a custom-made EMF stimulation device that encompasses the bioreactor and delivers uniform EMFs. We established a three-step EV purification protocol that enables high-density production of EVs. We then performed mass spectrometry-based proteomics analysis on EV-related proteins and used high-resolution nanoparticle flowcytometry for single-vesicle analysis. We demonstrate that electrical stimulations of current amplitudes at physiological level that are currently applied in therapeutic deep brain stimulation can modulate EV content in a frequency-dependent manner, which may have important implications for basic biology and medical applications. First, it raises intriguing questions about how the endogenous electrical activity of neuronal and other cellular assemblies influence the production and composition of EVs. Second, it reveals an additional underlying mechanism of how therapeutic electrical stimulations can modulate EVs and treat human brain disorders. Third, it provides a novel approach of utilizing electrical stimulations in generating specific EV cargos.

## Introduction

Electromagnetic fields (EMF) produced by moving electrical charges (*2*) with a wide frequency spectrum are a combination of invisible fields to visible light and beyond. The EMF has a vast range of everyday applications, from wireless cellular communications, electrical brain stimulation, to medical imaging, that rely on the fundamental aspects of the EMF frequency, which plays a critical role in applications from the behavior of subatomic particles to function of complex biological systems. Extracellular vesicles (EVs) are small membrane-bound bubbles released by living cells into the extracellular space and function as critical mediators for intercellular communications. EVs contain a wide range of bioactive molecules, including proteins, lipids, and nucleic acids, that can be transferred from one cell to another (*3*). Thus, using EVs as biomarkers or delivery vehicles for diagnosis and treatment is a rapidly growing field of research and clinical practice. Modulating the release, size distributions, and cargo content of EVs through EMF is an emerging research area that has many potential implications on both basic biology and therapeutic development. One of the most intriguing questions from this finding is how endogenous electrical activity emerging from neuronal assemblies, i.e., the local field potential, could affect the dynamics of cells releasing EVs and subsequently change biological functions. Secondly, this finding reveals additional mechanism for why therapeutic electrical brain stimulation are effective in treating human brain disorders or other diseases. Thirdly, it offers a novel method for producing pharmaceutical EVs, which is advantageous over other methods using chemical reagents. Electrical stimulation is a clean, controllable, and versatile method that can vary in frequency, field strength, and waveform morphology.

Four years ago, we postulated a hypothesis (*1*) suggesting that external EMF at physiological amplitudes would affect the release, size distribution, and content of EVs. In that manuscript we demonstrated how a low-frequency EMF altered the size distribution of EVs released from rat primary astrocytes. However, one of the major technical barriers was the lack of an ideal EMF stimulation device that could create a spatially uniform electric field. We subsequently published related work studying the frequency-dependent effect of EMF on the mobility of intracellular vesicles in astrocytes (*4*). During the last serval years, we made several technical improvements to the experimental protocol. First, we cultivated cells in a hollow-fiber bioreactor (Fibercell System) that provides *in vivo*-like environment and continuous EV productions over a longer period when electrical stimulation is applied; Second, we designed a novel stimulation device that encompasses the bioreactor and delivers uniformly distributed EMF over the bioreactor geometry, so that all collected EVs are released from cells under the same experimental condition (see Figure 1 A); Third, we developed a unique three-step purification method that greatly improves the purity and production of EVs, but avoids the use of ultracentrifugation method known to cause damages to EVs (*5, 6*). These advances can facilitate future mass production of EV cargos in uniform populations.

**Figure 1.**
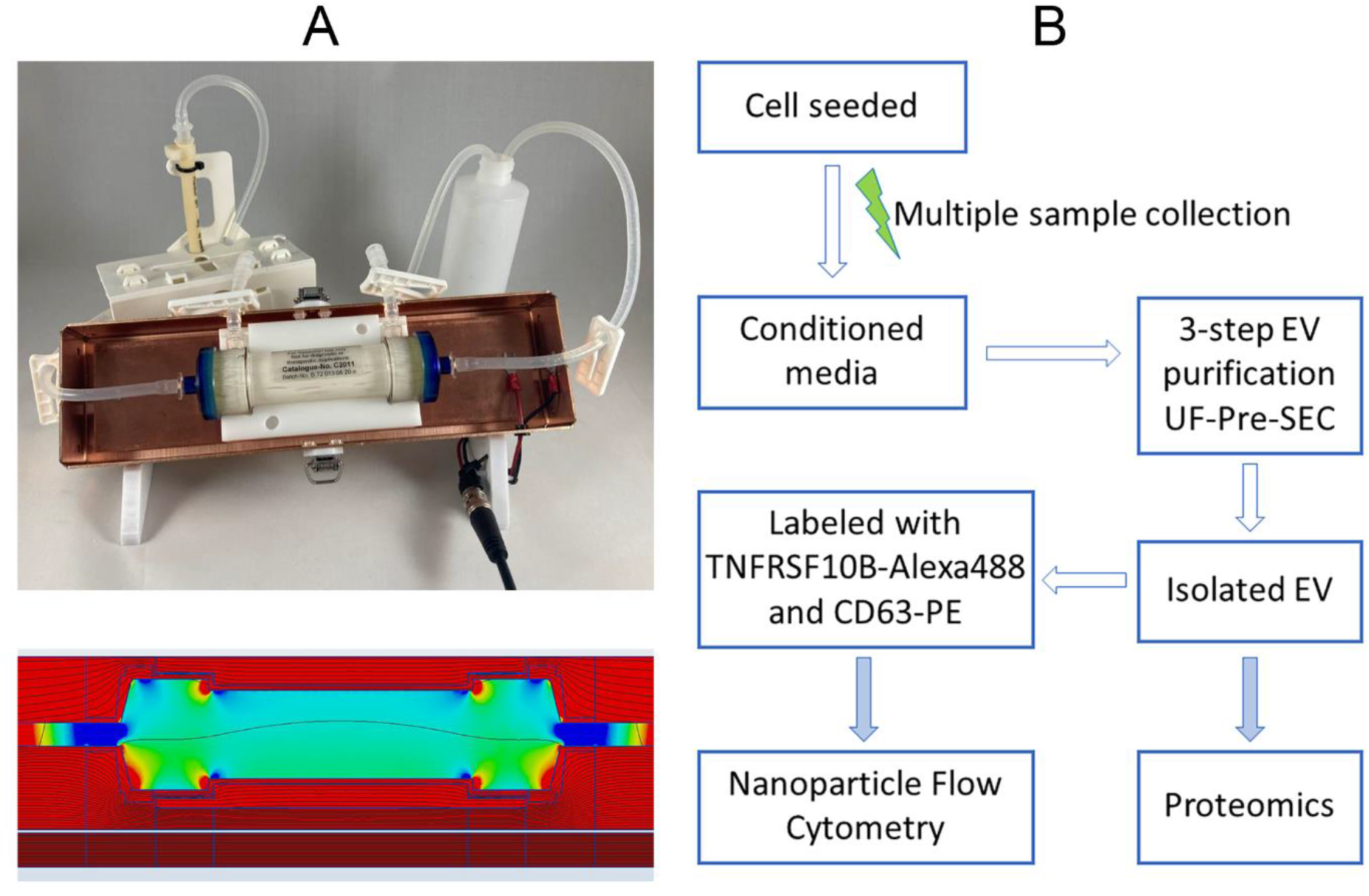
The pictures of Fibercell system and the experimental procedure flow chart The custom-made EMF stimulation device and a flowchart of experimental procedure. A) top: a Fibercell cartridge sit inside the stimulation device; bottom: a map showing the simulated electric field. B) The experimental procedure of EV purifications and detections.

To evaluate the stimulation device and examine the improved purification protocols, we choose a human cell line (HT-1080) over primary glial cells used previously (*1*), due to the requirement of seeding-cell number (> 100 million) for the medium-size Fibercell cartridge. The HT-1080 is a human fibrosarcoma cell line commonly used in cancer research and has been extensively characterized both genetically and phenotypically. These cells are known to be highly invasive and have been used as a model system to investigate the molecular mechanisms underlying cancer progression and test for potential cancer therapies. As proof-of-principal, we used HT-1080 as an example to demonstrate how EMF affects EVs in a frequency-dependent manner. EMF affecting EVs is likely a general phenomenon to all living cells, but different cell types could have their unique responses to a different spectrum of frequencies.

### Technical Innovations

1. A novel EMF stimulation device delivers uniformly and spatially distributed electrical fields for the Fibercell 3D culture system (details in a separate paper).
2. The 3-step EV-purification protocol is a unique combination of three methods including the ultrafiltration (UF), polyethylene glycol-based precipitation (Pre), and the size-exclusion chromatography (SEC) that circumvents weaknesses induced by individual method if used alone.

## Results

We performed three rounds of experiments in total, with each round comprised of four different conditions that include the control condition without electrical stimulation (No-ES) and three other conditions with electrical stimulation (ES) at different frequency of 2 Hz (low), 20 Hz (intermedia) and 200 Hz (high), respectively. For each condition, we cultivated HT1080 cells in a fresh Fibercell cartridge for a two-week period and harvested conditioned media five times (see methods for details). The EVs collected from the first round were used for protocol optimization, in which we added a membrane-permeable live-cell labeling dye (Calcein-AM) to distinguish intact EVs from other nano particles or soluble proteins (*7*). We used fluorescence-enhanced size-exclusion chromatography to confirm that the first elution peak of UV absorbance at 280 nm superimposes with the unique peak of the fluorescent signal for the EV population (*8*). This pilot experiment indicated that the first peak of UV signal can be used to determine the EV elution is clean and purified. Because the fluorescence from Calcein-AM could contaminate the mass spectrometry and nanoflow cytometry analyses, in the 2^nd^ and 3^rd^ rounds of EV collections we didn’t add Calcein-AM and only subjected them to downstream analysis following the flowchart illustrated in Figure 1B.

### 1. Proteomics study

We provided twenty-four samples of purified EVs (two rounds, four conditions with three consecutive samples per condition) to the Mayo Clinic Proteomics Center for mass spectrometry data analysis (see Methods for details). In the last batch of samples, a total number of 2,078 proteins was identified, 732 proteins were present in all samples, which were used in our analysis (see Table S1 in supplements).

To describe how EMF affects EV protein expression, we adopted the relative-difference (δ) algorithm (*9*) that calculates changes on consecutively collected samples within the same experimental condition (see Methods for details). A positive δ value means an increased protein expression while a negative δ value means a decreased protein expression under the corresponding condition.

We found that expressions of EV-related proteins varied with different EMF frequencies. Figure 2 is a heatmap summarizing of 732 proteins with the red color indicating the most positive δ value and the blue color showing the most negative δ value. At the bottom are four pie diagrams showing percentages for increased/decreased protein expression under four different experimental conditions. During cell cultivation inside the cell cartridge, protein expressions are variable even for the control condition. We found that 176 (24%) of 732 identified proteins had increased expression for No-ES; 214 (29%) for ES at 2 Hz, 158 (22%) for ES at 20 Hz and a sizable 422 (58%) for ES at 200 Hz. It is worth noting the significantly large percentage of increased protein expression under ES at 200 Hz.

**Figure 2.**
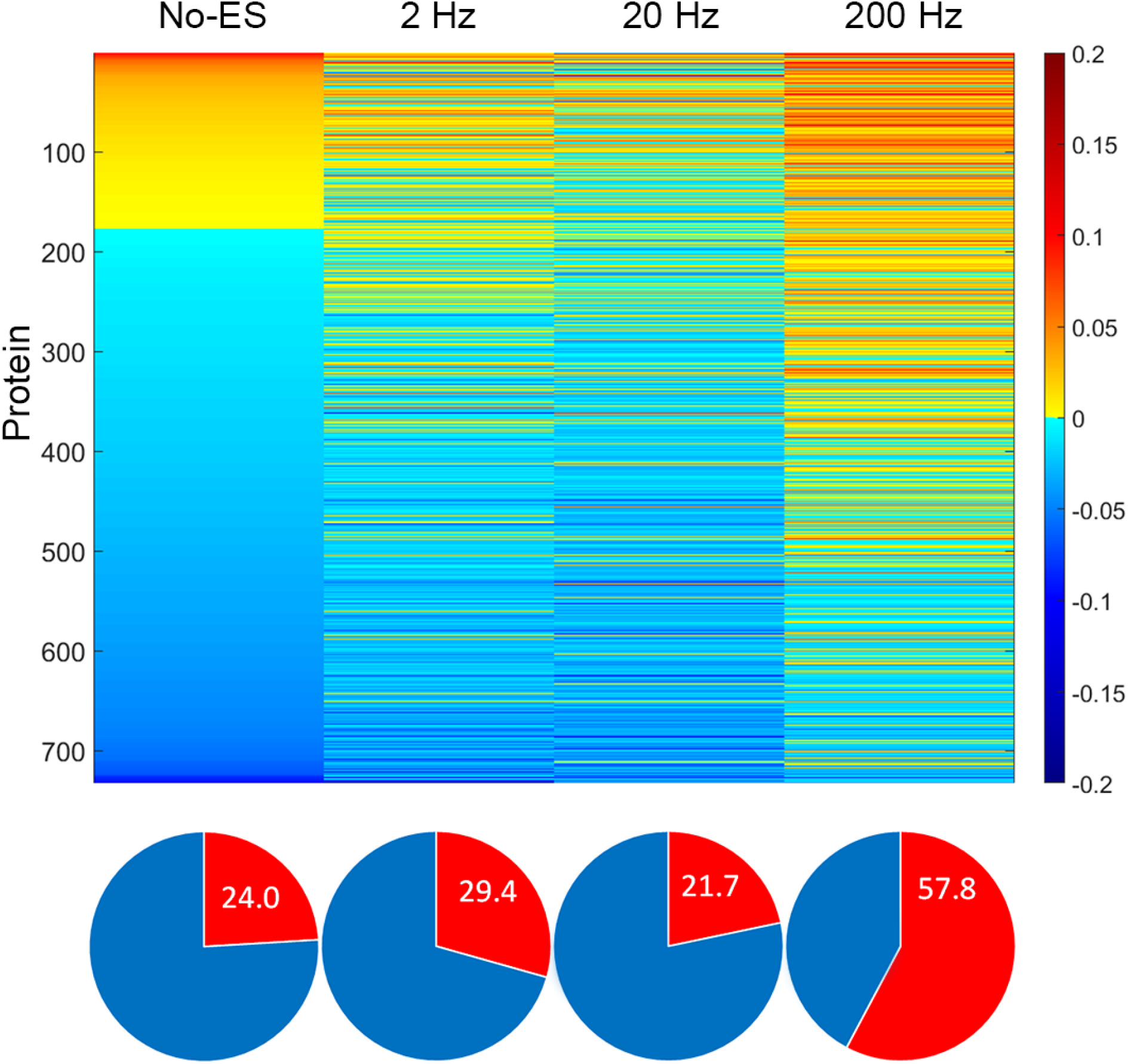
Heatmap and pie charts of 732 proteins A heatmap summarizing mass spectrometry analysis of 732 proteins. Each line in the heatmap represents the δ value of a unique protein, with its color reflecting the intensity labeled in the scalebar on the right that is proportional to the δ value. At the bottom are pie diagrams showing percentages of proteins with increased level (in red) under four different experimental conditions.

To gain an in-depth understanding of the biological meaning of these proteins, we performed enrichment analysis based on previously published results using DAVID (the database for annotation, visualization, and integrated discovery) (*10*) to extract biological features associated with the list of 732 protein genes. We found 115 types of biological processes associated with 450 genes (Table S2).

To bridge the connection between EMF effects and biological features, we used knowledge graph (KG) that connects extracted biological features to associated proteins along with their calculated δ values. However, the graph illustrating the entire network for 732 proteins is overwhelming for human eyes, we therefore created a compact list that only includes top ten candidates with the most positive δ values and ten others with the most negative δ values for each experimental condition. After taking out twenty-one duplicates, the final concise list contains 59 proteins (see Table S1-short in supplement), which was used to illustrate how EMF affects biological features of EVs. There were 37 types of biological processes associated with 34 proteins (Table S2-short). For visualization, we included those biological processes connected with two or more associated proteins. The resulting KG consisted of 17 biological processes and 29 proteins.

In Figure 3, we combined four KGs representing data from their corresponding experimental conditions, in which biological processes were illustrated with filled grey octagons and associate proteins were indicated by filled circles, with their radii proportional to corresponding absolute δ values. Blue/red color indicated genes for decreased/increased protein expression. Grey lines represented previously reported connections between biological processes and associated proteins.

**Figure 3.**
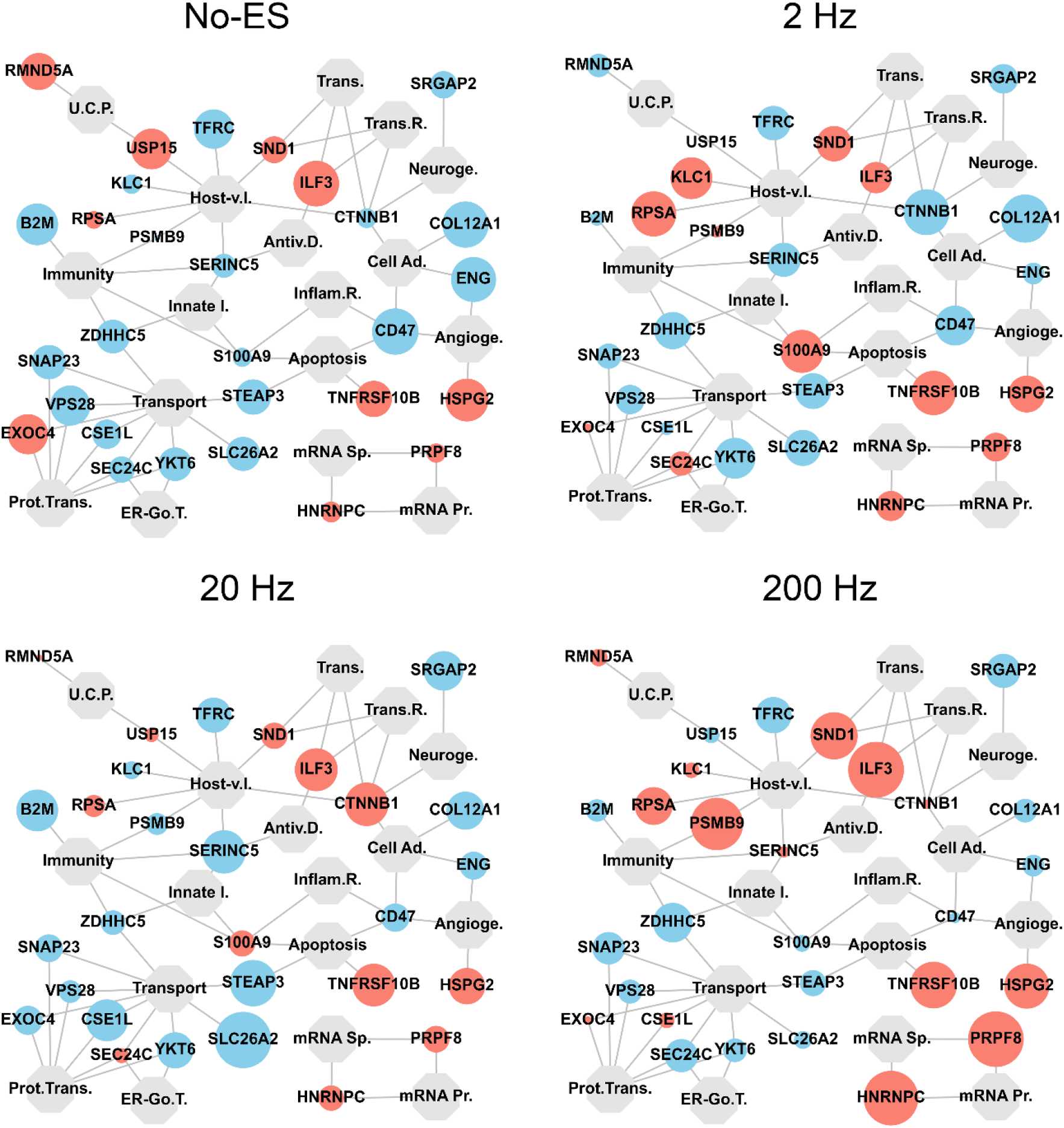
Knowledge graphs representing data from four different experimental conditions. Knowledge graphs showing biological processes associated with proteins. Grey octagons represent identified biological processes, whereas red/blue cycles indicate proteins with positive/negative δ values. The radius of each cycle is proportional to the absolute δ value of corresponding protein. Grey lines represented previously reported connections between biological processes and associated proteins. Abbreviations of biological processes: Host-v. I.: Host-virus interaction, Prot.Trans.: Protein transport, Angioge.: Angiogenesis, Cell Ad.: Cell adhesion, ER-Go.T.: ER-Golgi transport, Antiv.D.: Antiviral defense, Innate I.: Innate immunity, Inflam.R.: Inflammatory response, Neuroge.: Neurogenesis, mRNA Sp.: mRNA splicing, mRNA Pr.: mRNA processing, U.C.P.: Ubl conjugation pathway, Trans.R.: Transcription regulation, Trans.: Transcription.

The KG is a class of message passing neural networks commonly used for processing data that connects objects with edges. It has been used for deep learning in computing science that is beyond our focus for this work (*11*). But, even with visual of these graphs without a deep analysis we identified selected biological features that were altered under ES at a specific frequency. For example, at the lower-right corner of each graph is a subgroup network for mRNA processing/splicing associated with two proteins (PRPF8 and HNRNPC). We observed significant increase of both proteins under ES at 200 Hz, suggesting that mRNA activities can be enhanced by ES at 200 Hz. Another interesting spot is at the identified protein of S100A9 that has associations with four biological features including immunity, innate immunity, inflammatory response, and apoptosis. We observed significant increases of S100A9 under ES at 2 Hz. The third interesting spot is at the protein of TNFRSF10B that is associated with apoptosis. We detected its biggest increase under ES at 2 Hz, suggesting that immunity/apoptosis could be enhanced by ES at a low frequency.

The mass spectrometry data represents total exosomal proteins in regardless of whether the identified protein is from the cytosolic origin or others located on EV surface. Identifying exosomal surface proteins is important because they not only carry information on their tissues of origin but also play an important role as critical mediators for intercellular communications. Furthermore, to answer the question of whether a detected elevation in protein level was due to the increased number of EVs or the increased surface expression per EV, we performed high-resolution nanoparticle flowcytometry that enables detection of EVs at single-vesicle level.

### 2. Nanoparticle flowcytometry

For flowcytometry, we used a fluorescent-tagged antibody targeting specifically to the extramembrane domain of a surface protein. Although screening through all EV surface protein could be helpful, for this proof-of-principal study we focused on a couple of selected targets under following criteria. 1) it must be an exosomal surface protein; 2) it has a higher fraction among all detected proteins; 3) it is a well-studied protein with established important biological relevance; 4) it has a commercially available antibody suitable for flow cytometry. As a result, we choose the TNFRSF10B (Tumor Necrosis Factor receptor superfamily member 10b, also known as the death receptor 5, DR5, or the TRAIL receptor 2, TRAIL-R2) as the evaluation target. TNFRSF10B contains an intracellular death domain, which can be activated by TNF-related apoptosis inducing ligands and transduces an apoptosis signal (*12*). Monoclonal antibodies targeting TNFRSF10B have been developed and are currently under clinical trials (such as Contumumab, Lexatumumab, Tigatuzumab and Drozitumab) for patients suffering from a variety of cancer types (*13-15*). Our question here is whether TNFRSF10B+ EVs are modulated by external EMF. We also used one of the common exosomal marker, CD63, as another targeting protein (*16*). An Alexa488-tagged TNFRSF10B antibody and a Phycoerythrin (PE)-tagged CD63 antibody were used for duo-labeling single-vesicle characterization using a nanoflow cytometry analyzer (FCM N30).

Figure 4A. includes scatter plots showing populations of EVs detected from flowcytometry with the excitation wavelength at 488 nm. The Alexa488 signal was represented by the vertical axis, whereas the PE signal was indicated in horizontal axis. For each scatter plot, the distribution was convened into four quarters based on baseline cutoffs on the green or red fluorescent signals. On the top-left corner (Q1) was the population for TNFRSF10B+ only EVs. Following clockwise on the right-top (Q2) is the population of EVs with duo-labeling of TNFRSF10B+/CD63+ EVs, and on the lower right (Q3) is the population for CD63+ only EVs. Q4 on the lower left is the distribution of non-specific EVs or other nano particles that are not considered further.

**Figure 4.**
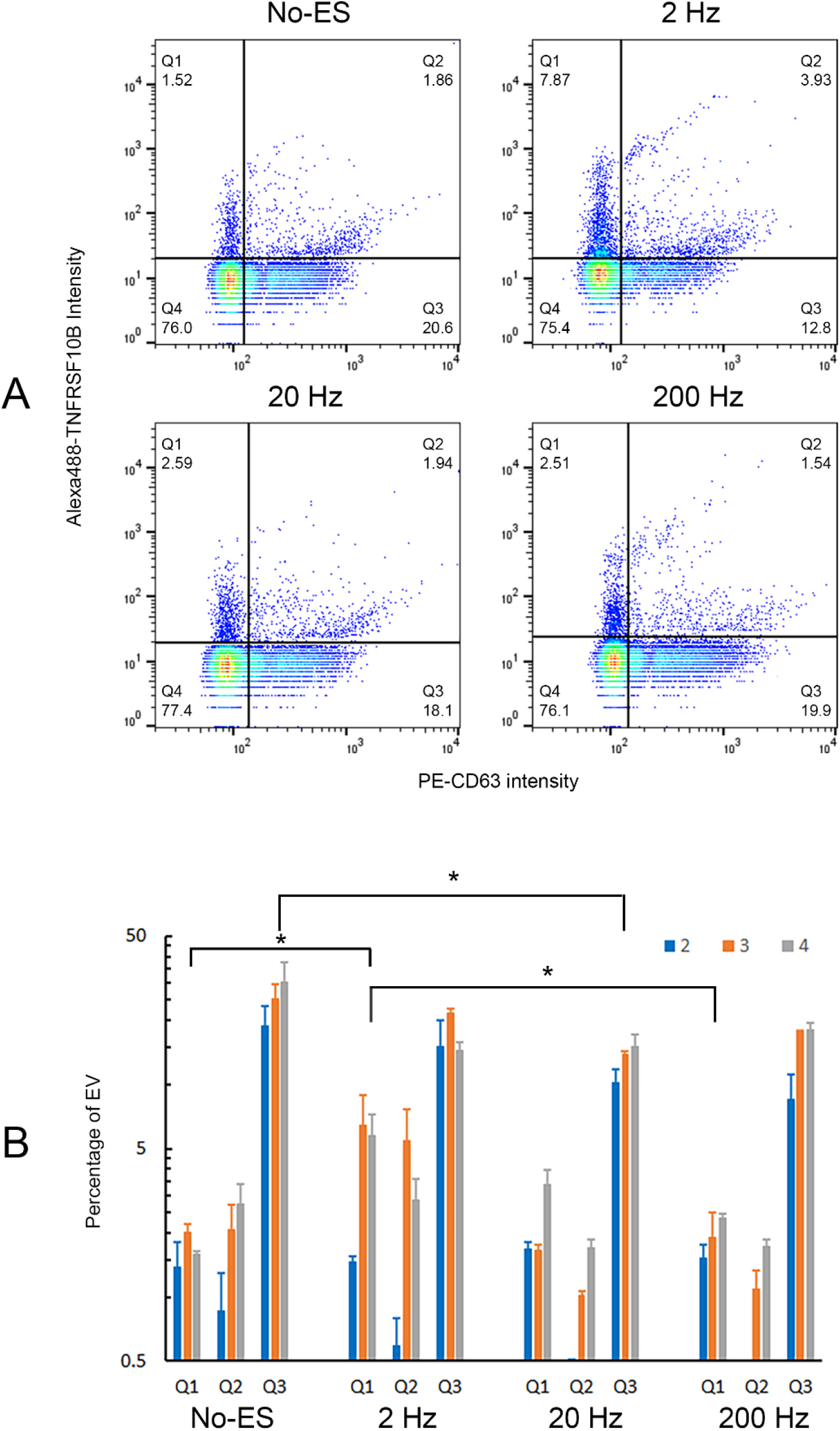
Scatter plots and bar graph of EVs detected from flowcytometry. Results from nano-particle flowcytometry detections of TNFRSF10B/CD63 dual-labeling EVs. A) Scatter plots display populations of EVs at four different experimental conditions. B) A bar graph compares each EV population, with the height of each bar indicates the percentage for EV population (P_Q1_, P_Q2_ and P_Q3_). Samples from the 2^nd^, 3^rd^ and 4^th^ collections were indicated in blue, orange, and grey colors, accordingly. (p < 0.05)

To illustrate changes of EVs, we used a bar graph (Figure 4B), in which the height of each bar indicates the percentage (P) for each population (P_Q1_, P_Q2_ and P_Q3_). Samples from consecutive collections (2^nd^, 3^rd^, and 4^th^) were labeled in blue, orange, and grey colors, accordingly. Four different experimental conditions were marked as No-ES, ES at 2 Hz, 20 Hz and 200 Hz, respectively.

For Q1(TNFRSF10B+ only), the P_Q1_ from 2^nd^ collections of all four conditions were around 1.5∼2 %. However, in the 3^rd^ and 4^th^ collections P_Q1_ under ES at 2 Hz were at ∼7-8% (about 4-fold increase) when compared to that of No-ES or ES at 200 Hz (p < 0.05). P_Q2_ under ES at 2 Hz had a similar increase from 2^nd^ to 3^rd^ or 4^th^ collections. Interestingly, P_Q3_ under ES at 20 Hz was lower than any other three conditions (p < 0.05). In conclusion, ES at 2 Hz frequency selectively increased the percentage of TNFRSF10B+ EVs, but ES at 20 Hz frequency reduced percentage of CD63+ EVs. But we did not observer significant changes for TNFRSF10B+ EVs under ES at 200 Hz.

We also measured the average size (Ø) for each EV population. The TNFRSF10B+/CD63+ dual-labeled population (Q2) had the largest size (> 90 nm), followed by the TNFRSF10B+ population (Q1) and the CD63+ population (Q3). But no detectable difference was found for the average size of individual population when comparing that from No-ES to ES at any of three frequencies (see Table 1), suggesting that EMF may not affect EVs-releasing mechanism but could modulate specific protein synthesis and packing along the EV-forming pathway.

**Table 1.**

| Table 1 | $\phi_{Q1}(\text{nm})$ | $\phi_{Q2}(\text{nm})$ | $\phi_{Q3}(\text{nm})$ | $\phi_{Q4}(\text{nm})$ |
| --- | --- | --- | --- | --- |
| No-ES | $84.2 \pm 1.6$ | $92.3 \pm 3.6$ | $76.0 \pm 3.7$ | $76.2 \pm 4.0$ |
| 2 Hz | $87.7 \pm 10.4$ | $93.6 \pm 8.0$ | $72.8 \pm 2.5$ | $74.0 \pm 2.9$ |
| 20 Hz | $84.4 \pm 6.2$ | $98.5 \pm 6.8$ | $75.6 \pm 3.6$ | $76.2 \pm 2.7$ |
| 200 Hz | $81.9 \pm 0.9$ | $93.4 \pm 3.5$ | $73.3 \pm 2.5$ | $74.2 \pm 2.0$ |

In summary, we performed mass spectrometry-based proteomics and nanoflow cytometry-based single-vesicle detections to investigate whether EMF affects EVs at a frequency-dependent manner. The gathered evidence shows that EMF at a lower frequency increases the percentage of EVs carrying specific exosomal proteins linked to the group of biological processes including immunity, inflammation, and apoptosis.

## Discussion

This is a continuation and expansion of work we started four years ago when we only released a preprint (*1*) due to the lack of a controllable EMF stimulation device. In the last several years we developed a custom-made electrical stimulation device that can deliver uniformly distributed electric field to a three-dimensional space where cells grow in a near-vivo environment. In a separate method paper we will describe the engineering and phantom EMF measurements of the device in detail. For this report, our discussions are mainly focused on three aspects: the unique 3-step EV-purification protocol; profile of exosomal protein; and the potential benefit of using EMF in cancer treatments.

### 3-step EV-purification protocol

One of the most challenging aspects, and often underappreciated, of EV research is to isolate EVs and accurately determine the exact exosomal protein composition with a minimum level of other soluble molecules or large protein complexes. Although ultracentrifugation (UC) is still the commonly used method for EV isolation, the yield of this procedure is low, and it does not guarantee a high purity (*5*). Other methods such as ultrafiltration (UF) and polymer-based precipitation (Pre) are less labor-intensive with low-speed centrifugation and easy in handling of large volume samples. But the purity of EVs from these methods is compromised due to the co-isolation of protein aggregates and abundant non-vesicular proteins (*17*). On the other hand, size-exclusion chromatography (SEC) separates large particles such as EVs from small particles represented as soluble proteins, but its small starting sample volume is a significant limitation.

Several recently published articles suggest a mixed use of different methods. Tethering UF with SEC can result in an overload of protein, which also undermines EV separations. The Pre-SEC method adds the precipitation step before loading the sample to the chromatography column to eliminate most soluble protein and increased the resolution of the SEC method (*17*). The Pre-SEC strategy was successful to conditioned media based on the low-glucose Dulbecco’s Modified Eagle Medium, but it has the precipitation issue the high-glucose medium supplemented with CDM-HD (Chemically Defined Media - High Density), a universal serum substitutes specifically designed for cells grown in hollow fiber bioreactor systems. To circumvent this issue, we added a UF step before Pre to minimize the number of glucose and other small molecules that may interfere with precipitations, and we used a lower KD cutoff UF filter (< 10 KD) to maximize recovery of EVs. This 3-step UF-PRE-SEC purification protocol is unique to high-density cell culture with CDM-HD. It is worthy to note that a recently available instrument named EXODUS is a one-step automatic EV purification machine aiming at the simplification for EV isolation (*18*). At the time of finishing this report, we didn’t have the opportunity to evaluate this instrument, so we cannot make any comments on how to compare between our method with that from EXODUS.

### Proteomic Profiles

We choose the Fibercell system (Fibercell Inc.) as a platform to investigate how EMF modulates EVs. Previous reports have shown that EVs generated with the hollow-fiber bioreactor technology have a low immunogenicity and immuno-regulatory antigenic signature (*19*). EVs can be produced in a large-scale system over a long production period that allows frequent harvest with the same quality and quantity (*19*). Therefore, our methods revealed here can be easily adopted to future large-scale production of EVs.

The selection of the two-week period for cell cultivation is based on a consideration for multiple sample harvestings needed for downstream analysis. Instead of comparing single data between different experimental conditions, we used the relative-difference algorithm that computes data from consecutive samples under the same experimental condition, so that any unknown issues caused by differences in seeding cells or by different cartridges can be kept at the minimum. The calculated relative differences of identified proteins reflect how EMF affects EVs over time.

We used KG to illustrate how EMF affects biological features with EVs as mediators and found that different biological features respond to different frequencies. For example, we found significantly elevated percentage of increased protein expression at 200 Hz frequency. Specifically, two proteins (PRPF8 and HNRNPC) associated with mRNA processing/splicing were elevated at 200 Hz, whereas the protein of S1009A associated with immunity/apoptosis was greatly increased at 2 Hz.

There is a significant literature on interactions between EMF and biological entities owning to the long history beginning from the late eighteenth century when Luigi Galvani first recorded the phenomenon while dissecting a frog and conducting experiments with static electricity (*20*). Importantly, there are recently published literatures coincided with our findings. For example, the extremely low frequency stimulation of electromagnetic fields can exert a certain action on autoimmunity and immune cells (*21, 22*), and cause apoptosis (*23*). Others reported that electrical stimulations of frequency from 100 Hz to 1K Hz increased protein synthesis in articular cartilage explants (*24*), which agree with our findings of increased mRNA processing at 200 Hz when considering that mRNA translation is a key focal point of gene expression regulation. It is worthy to note that most published literatures were on interactions between EMF and tissues or cells. Our report was the first to our knowledge that extends EMF effects to EVs.

EVs also carry different nucleic acids, including microRNAs that can significantly regulate cell growth and metabolism by post-transcriptional inhibition of gene expression. Examine the microRNA profile will not only allow the determination of the status of the cells and their origin but will allow analyses to their targeted cell groups and possible functions. Studying microRNA profile is another important aspect in our future direction of high-throughput genomics study of EVs.

### Nanoflow Cytometry

Different from the method used in mass spectrometry studies, plasma membranes of EVs were kept intact during sample preparation for nanoflow cytometry. Without a membrane-permeation reagent, fluorescence-tagged antibodies were limited on accessing EV surface proteins, whereas proteins from the cytosolic origin were not detected. We used antibodies targeting at the extramembrane domain of TNFRSF10B and CD63 for dual-labeling experiments and found three distinguished EV populations, including the TNFRSF10B+ only, the CD63+ only and the TNFRSF10B+/CD63+ dual-labeled subgroups.

These results suggested the complexity of EVs and reminded us that we should be cautious when using a commonly recognized EV makers (like CD9, CD63 or CD81) to describe all EVs. Here, CD63 represents only a certain percentage among all EVs. Similarly, TNFRSF10B has an even lower appearance. Clearly, the subgroup of TNFRSF10B+ only EVs is very different from the subgroup of CD63+ only EVs, as indicated by their size measurements through side scattered light. Among three EV subgroups, the TNFRSF10B+/CD63+ dual-labeled subgroup has the largest average size, followed by the TNFRSF10B-only labeled subgroup and then the CD63-only subgroup (Table 1). There is a possibility that the measured average size could be artificially expanded due to attached antibodies when the size of a vesicle is on the similar scale to the antibody-tag complex. Single vesicle with more attached antibodies would appears to be in a larger dimension based on the side scattered light detection methodology. Alternatively, using Cryo-EM technology to detect the EV size would be the optimal tool for size determinations of EVs, which we will consider in future investigations. Nevertheless, we didn’t observe any EMF induced change in the average size of an individual subgroup. A similar observation (unpublished) was concluded by Dr. Miscenko when he presented at the recent ISEV2023 meeting.

Interestingly, we found 3∼4-fold increasing of TNFRSF10B+ subgroups at a low frequency of 2 Hz. TNFRSF10B is mainly located on plasma membrane. As an important mediator of the extrinsic pathways of apoptosis, TNFRSF10B has quickly emerged as the target of therapies for cancer treatments that utilize two type of pharmaceutical agents, recombinant human TRAIL proteins (such as Dulanermin) and TRAIL-R2 agonist antibodies (such as Conatumumab, Lexatumumab, Tigatuzumab and Drozitumab) (*13-15*). Although the preclinical results for TRAIL-R2 agonist antibodies were promising, the response/recovery rate was low when they were tested in patients (*13, 25*). A recent study suggested that the abundance of TNFRSF10B on surface of cell could be regulated by vesicle transport and the higher expression of TNFRSF10B were correlated with higher first/post-progression survival in chemotherapy-treated lung cancer patients (*26*). Our results of increased population of TNFRSF10B+ EVs at a low frequency of EMF suggested a potential therapeutic treatment through the combination of chemotherapy with EMF enhancement that would be beneficial to patients who have decreased response to initial TRAIL-R2 treatment.

Frequencies of endogenous EMF in certain body regions are varying through regular activities and are associated with physiological conditions. For example, a recent study showed that patients with Parkinson’s disease have different EEG frequency pattern in the supplementary motor area, accompanied by reduced walking speed and step length. When patients walked with enhanced arm swing, their EEG frequency pattern, walking speed and step length became normal (*27*). On the other hand, accelerating evidence showed that the frequency of EMF stimulation has dominant effects. For example, one hour of ES at 20 Hz dramatically accelerates axonal regeneration in rat (*28*). ES at 2 Hz achieved the best effect for promoting proliferation of satellite cell in muscles disuse atrophy (*29*).

However, most previously reported studies of EMF effects on biological functions were focused on the field intensity rather than the frequency (*30, 31*), and the applied intensity was also much high than endogenous electrical field. As of today, we know little about the mechanistic explanation of how endogenous electrical activities affect biological functions. Our findings from this report could be the key to uncover the real functionality of endogenous electrical activities.

We all inhabit a universe charged with EMF, either originating from the deep cosmos or created by mankind. Like the symphony orchestra, both endogenous and exogenous electrical activities are in harmony, inspiring people to reach, strive and dream through a unique combination of frequencies. To conclude this discussion, we would like to share a famous quote by Nicola Tesla: “If you want to find the secrets of the universe, think in terms of energy, frequency and vibration”.

## Methods

### Cell culture

Human fibrosarcoma cell line (HT1080) was cultivated following the standard protocol with DMEM at 37° C with 5% CO_2_. The low glucose culture media included DMEM (1 g/L glucose, Gibco 11885-084, Thermo Fisher Scientific, Waltham, MA) supplemented with 10% (v/v) Fetal Bovine Serum (FBS, Gibco 10437-028, Thermo Fisher Scientific) and 1% (v/v) Penicillin Streptomycin (Gibco 15140122, Thermo Fisher Scientific). The high glucose culture media included DMEM (4.5 g/L glucose, Gibco 11965-092, Thermo Fisher Scientific), 10% (v/v) CDM-HD serum replacement (FiberCell Systems Inc., New Market, MD) and 1% (v/v) Penicillin Streptomycin (Gibco 15140122).

### Generation of cell conditioned media

We used a 3D cell culture system from FiberCell for all experiments. Each fresh cell cartridge (C2011, FiberCell Systems Inc.) was prepared according to the manufactory guideline through sequentially wash with PBS for 72 hours, DMEM + 1% Penicillin-Streptomycin for 28 hours and DMEM+10% FBS+1% Penicillin-Streptomycin for 44 hours. Normally, each cell cartridge allows up to 100 days for conditional media collections. We limited our collection duration to 2 weeks. To maximize the efficiency of each cartridge, we double the required cell seeding number and started with 2 × 10^8^ cells on day 1 for each experiment. Five sequential cell conditioned media was then collected on day 3, 5, 8, 10 and 12 respectively. Electrical stimulations were started on day 3 right after the first conditional media was collected and stopped till day 12 after the fifth sample collection. The low glucose culture media was replaced by the high glucose culture media after the second sample was collected on day 5. The glucose level of the cell culture media was monitored daily using blood glucose meters (AimStrip Plus, Germaine laboratories, INC, San Antonio, TX; ACCU-CHEK Guide Me, Roche Diabetes Care, Inc., Indianapolis, IN). We monitored temperature on the surface of cartridge in all experiment using a thermometer (TENMA 72-7715, Newark, Chicago, IL). Measured temperatures were maintained at 37.0±0.2 °C.

### Electrical stimulation

The electrical stimulation was applied through a custom-designed device that delivers a uniformly distributed electromagnetic field with a controlled frequency and amplitude. EMF signals were created with a function generator (4055B, B&K Precision, Yorba Linda, CA). We applied a square-wave pulse with 0.4 ms in width and 4 V amplitude that produces an equivalent electrical field strength of 5 mV/mm within the cartridge space according to a simulation performed with the QuickField 6.4 software (Tera Analysis Ltd, Columbia SC, USA). All stimulation protocols follow a repeated 2-min pattern that includes 120 pulses. For ES at 2 Hz, all 120 pulses were evenly delivered within the first minute and leave the second minute being quiet. For ES at 20 Hz, 120 pulses were delivered within the first 6 seconds and leave the rest 114 seconds being quiet. Similarly, for ES at 200 Hz, 120 pulses were delivered within the first 0.6 seconds, and leave the rest 119.4 seconds being quiet. So that, the total number of delivered pulses for each ES conditions were equal. Quiet periods were necessary to avoid over heating caused by applied electrical field.

### EV purifications

20 mL of cell conditioned media was first subjected to a low-speed centrifuged at 3000 × g (20 min, 4°C) to remove cell debris, with the supernatant filtered through a 0.22 μm syringe filter (Millex-GS, SLGS033SS, MilliporeSigma, Burlington, MA). Then we followed the 3-step purification protocol: 1) reducing sample volume using 10 KD Amicon™ Ultra-15 centrifugal filter units (UFC 901024, MilliporeSigma™); 2) removing mostly soluble proteins with self-prepared polyethylene glycol precipitation (PEG) solution. In details, a working PEG solution was prepared with PEG8000 (Sigma Aldrich) at 24% with PBS and 75 mM NaCl. Concentrated cell conditioned media from step 1 was thoroughly mixed with the working PEG solution at 2:1 volume ratio (final concentration of 8% PEG), stored at 4°C overnight and finally centrifuged at 1,500 × g for 30 minutes at 4°C. The obtained pellet was dissolved in 500 μL of calcium/magnesium-free PBS; 3) removing free polyethylene glycol and other remaining soluble proteins using size-exclusion chromatograph (SEC). In detail, 500 μL sample from step 2 was overlaid to a self-made SEC column (Bio-Rad Glass Econo-Column Chromatography Columns, 7371512, packed with 10 mL Sepharose CL-6B, 17016001, from Cytiva, Muskegon, MI). A low-pressure liquid chromatograph system (Biologic LP, Bio-Rad) was used to deliver a constant flowrate at 500 μL/min with the collecting fraction volume of 200 μL. Purified EVs were recovered from the first elution peak that includes 5 fractions with most protein concentrations measured by the UV absorbance at 280nm. We took 40 μL from each of the 5 fractions, pooled together, and stored at -80 °C till proteomics analysis. Remaining purified EV samples were stored at 4 °C till nanoparticle flow cytometry analysis.

### Nanoparticle flow cytometry analysis

High-resolution flowcytometry for nanoparticle analyses were performed using the Flow Nanoanalyzer (N30, NanoFCM, Xiamen, China). In details, 100 μL of isolated EV sample was mixed with 4 μL of Alexa488 anti-human TNFRSF10B antibody (FAB6311G, R&D Systems, Minneapolis, MN) and 2 μL of PE anti-human CD63 antibody (561925, BD Biosciences, Franklin Lakes, NJ) and incubated at 37 °C for 30 minutes. Unlabeled antibodies were removed using second SEC with the qEVsingle column (70 nm Gen 2, IZON, Medford, MA). Again, labeled EVs were recovered in the first elution peak and then subsequently subjected to nanoparticle data collections. All data were converted to FCS 3.0 using NF Profession (Version 1.0, NanoFCM) and analyzed with FlowJo (Version 10.8.1, BD, Franklin Lakes, NJ).

### Mass spectrometry data acquisition and processing

Mass spectrometry (MS) based proteomics of EV-related proteins were performed at the Mayo Clinic Proteomics Core. In details, 200 μL of each isolated EV sample were first precipitated with cold acetone, then digested with trypsin and extracted using the S-Trap micro kit (ProtiFi, Fairport, NY). The concentration of peptide was determined using Pierce quantitative fluorometric peptide assay (Cat# 23290, Thermo Scientific, Rockford, IL). 11 – 18 μL of the peptides were analyzed through nano-ESI-LC/MS/MS using the Orbitrap Exploris mass spectrometer coupled to a Dionex nano-LC system (Thermo Scientific, Waltham, MA). Liquid chromatography (LC) was performed using multistep linear gradients with solvent A (2% acetonitrile and 0.2% formic acid in H_2_O) and solvent B (80% acetonitrile, 10% isopropyl alcohol, and 0.2% formic acid in H_2_O) under a constant flow rate of 300 nL/min. The mass spectrometer was set at a resolution of 60,000 (at 200 m/z) in data dependent acquisition, with a full MS1 scan ranging from 340 to 1600 m/z. Dynamic exclusion was set to 25 seconds with cycle time in 3 seconds.

All raw MS files were analyzed using MaxQuant (Version 1.6.17.0) (*32*). MS/MS spectra was set up to search against the SwissProt (*33*) Human Database (2022 ver. 1), assuming trypsin digestion with up to two missed cleavages with the fragment ion tolerance of 20 PPM and parent ion tolerance of 4.5 ppm. Cysteine carbamidomethylation was set as a fixed modification and methionine oxidation was set as a variable modification. The false discovery rate was set to 0.01 for protein level and peptide spectrum match. Protein identification required at least one unique or razor peptide per protein group. Only proteins with unique peptide were used in further data analysis. Contaminants, and reverse identification were excluded from further data analysis.

### Bioinformatics analyses

We used relative intensity based absolute quantification (riBAQ) to determine the relative molar abundances of proteins, which normalized each protein’s iBAQ value to the sum of all iBAQ values from that sample (*34*). The heatmap was generated using MATLAB (The MathWorks Inc (2020). MATLAB Version: 9.8.0 (R2020a)). We used the relative difference to show effects of ES on the abundance of EV-related proteins. Calculations of relative differences (δ) for each identified protein (k) were performed using the following equation.

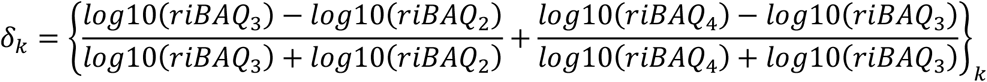

For each experimental condition, we discarded the first and the last samples and only used the middle three collected samples, namely 2, 3, and 4 representing samples collected on day 5, day 8 and day 10, respectively. The y-axis in the heatmap represents δ values. To determine whether the protein has been identified as a membrane protein or existing in extracellular exosome, we used a custom code in MATLAB to search against the SwissProt human database (downloaded in March 2023, 69670 entries; 2118 entries). The biological processes were obtained using DAVID bioinformatics resource (*35, 36*). Knowledge graphs were generated by the Python language package of NetworkX (*37*).

## Statistical evaluations

To determine the significance of differences in the percentage of TNFRSF10B/CD63 positive EVs between different experimental conditions, we performed a two-sample weighted Kolmogorov-Smirnov test (KS-test) using the Python package of SciPy (*38, 39*). The weight was determined using exponential smoothing methods (*40*). Other codes were all written in MATLAB.

## Supporting information

Supplemental Table S1-short

Supplemental Table S2-short

## Acknowledgement

We acknowledge the assistance of the Mayo Clinic Proteomics Core, which is a shared resource of the Mayo Clinic Cancer Center (NCI P30 CA15083). This work was supported by an NIH BRAIN initiative R01 grant to HLW and GAW (NS112144 from NINDS and NIMH) and the Minnesota Partnership for Biotechnology and Medical Genomics (MNP #17.16).

## Author Contributions

H. W. designed experiments; H.W. and Y.W. carried out the experiments and analyses; Y.W., H.W. and G.W. wrote the paper.

## Conflict-of-interest statement

The authors have no conflicts of interest to declare. All co-authors have seen and agree with the contents of the manuscript and there is no financial interest to report. We certify that the submission is original work and is not under review at any other publication

## Notes

### Competing Interest Statement

The authors have declared no competing interest.

